# Nanopore sequencing reveals full-length Tropomyosin 1 isoforms and their regulation by RNA binding proteins during rat heart development

**DOI:** 10.1101/2020.07.30.229351

**Authors:** Jun Cao, Andrew L. Routh, Muge N. Kuyumcu-Martinez

**Author notes:** Cardiology Department, Boston Children’s Hospital, Harvard Medical School, Boston, MA 02115. Corresponding author: Muge N. Kuyumcu-Martinez.

## Abstract

Alternative splicing (AS) contributes to the diversity of the proteome by producing multiple isoforms from a single gene. Although short-read RNA sequencing methods have been the gold standard for determining AS patterns of genes, they have a difficulty in defining full length mRNA isoforms assembled using different exon combinations. Tropomyosin 1 (TPM1) is an actin binding protein required for cytoskeletal functions in non-muscle cells and for contraction in muscle cells. *Tpm1* undergoes AS regulation to generate muscle versus non-muscle TPM1 protein isoforms with distinct physiological functions. It is unclear which full length *Tpm1* isoforms are produced via AS and how they are regulated during heart development. To address these, we utilized nanopore long-read cDNA sequencing without gene-specific PCR amplification. In rat hearts, we identified full length *Tpm1* isoforms composed of distinct exons with specific exon linkages. We showed that *Tpm1* undergoes AS transitions during embryonic heart development such that muscle-specific exons are connected together generating predominantly muscle specific *Tpm1* isoforms in adult hearts. We found that the RNA binding protein RBFOX2 controls AS of rat *Tpm1* exon 6a, which is important for cooperative actin binding. Furthermore, RBFOX2 regulates *Tpm1* AS of exon 6a antagonistically to the RNA binding protein PTBP1. In sum, we defined full length *Tpm1* isoforms with different exon combinations that are tightly regulated during cardiac development and provided insights into regulation of *Tpm1* AS by RNA binding proteins. Our results demonstrate that nanopore sequencing is an excellent tool to determine fulllength AS variants of muscle enriched genes.

## INTRODUCTION

Gene regulation by alternative splicing (AS) is an important contributor to development and tissue identity (Baralle and Giudice 2017). AS not only controls gene expression but also generates different isoforms of genes. Genome-wide analyses indicate that the majority of human genes undergo AS (Wang et al. 2008). Currently, many computational approaches based on short read RNA-sequencing are available to investigate AS patterns. However, with these techniques it can be difficult to determine the connectivity of multiple exons in a given transcript. This becomes more challenging if a given gene has many potential isoforms. Recent advances in nanopore sequencing technology allow sequencing of ultra-long DNA sequences (Lu et al. 2016) (Jain et al. 2018). The Oxford MinION sequencer is a portable device that provides realtime, high-throughput and long-read sequencing with <10% error rate (de Jong et al. 2017; Oikonomopoulos et al. 2016; Wang et al. 2020). This technology, therefore, is very attractive to study complex AS patterns in the context of full-length transcripts.

TPM1 is a coiled-coil protein that wraps around the actin molecules and provides stability to actin filaments. TPM1 is the predominant tropomyosin gene expressed in cardiac muscle and plays a significant role in muscle contraction (Bai et al. 2013). TPM1 is required for myofibril organization (Thomas et al. 2010), myocardial contraction (Wolska and Wieczorek 2003), and cardiac development (England et al. 2017). Mutations or aberrant expression of *TPM1* are associated with familial hypertrophic cardiomyopathy (Jongbloed et al. 2003; Marques and de Oliveira 2016; Muthuchamy et al. 1999; Thierfelder et al. 1994), dilated cardiomyopathy (Karam et al. 2011; Redwood and Robinson 2013) and heart failure (Rajan et al. 2010). *TPM1* has 15 exons, several of which are alternatively spliced, generating many gene isoforms generated via AS that are tissue specific and developmentally regulated (Gooding et al. 2013a). These multiple isoforms render distinct functions including cytoskeleton support and muscle contraction in the heart (Gunning et al. 2015; Lin et al. 2008; Perry 2001; Schevzov et al. 2011). However, it is still unclear what mechanisms dictate highly coordinated AS of *Tpm1* that impacts its expression and function in a cell- and development-specific manner. In this study, we used nanopore sequencing to identify full length transcripts of *Tpm1* gene with complex AS patterns in rat hearts at different developmental stages.

RBFOX2 is an RNA binding protein, which regulates AS by binding to a highly conserved motif ((U)GCAUG) in pre-mRNAs (Huang et al. 2012; Lovci et al. 2013; Sun et al. 2012; Yeo et al. 2009). RBFOX2 is important for muscle differentiation (Sebastian et al. 2013), maintaining muscle mass (Singh et al. 2018a), and sustaining muscle function (Gallagher et al. 2011). We and other groups have shown that RBFOX2 is involved in cardiovascular diseases including hypoplastic left heart syndrome (Homsy et al. 2015; McKean et al. 2016; Verma et al. 2016), heart failure (Wei et al. 2015), and diabetic cardiomyopathy (Nutter et al. 2017; Nutter et al. 2016). RBFOX2 binding sites are enriched near alternative exons that are developmentally regulated postnatally in the heart (Misra et al. 2020), suggesting a role for RBFOX2 in regulation of AS during heart development.

Using nanopore cDNA sequencing, we identified full length *Tpm1* isoforms with unique exon combinations that are regulated during rat heart development. We found that muscle and non-striated muscle *Tpm1* isoforms was generated via AS of specific exons during rat heart development. We uncovered that RBFOX2 regulates AS of rat *Tpm1* exon 6a. Furthermore, we found that RBFOX2 and PTBP1 antagonistically control AS of *Tpm1* exon6a. Overall, our results reveal that *Tpm1* spliced isoforms are tightly regulated during rat cardiac development and that RNA binding proteins RBFOX2 and PTBP1 have opposing roles in controlling developmentally regulated *Tpm1* AS. Our findings have broad implications in defining complex alternative splicing patterns of abundant cardiac muscle-enriched genes using nanopore cDNA sequencing.

## RESULTS

### Nanopore sequencing identifies full length *Tpm1* isoforms that undergo alternative splicing transitions during rat heart development

*Tpm1* has many different isoforms generated via AS (Gooding et al. 2013a). To determine the exact combination of exons in full length *Tpm1* isoforms in the heart at different developmental stages, we used total RNA from embryonic day 20 (E20, n=3) and 6-month (6M, n=3) rat hearts and generated cDNA for nanopore sequencing on the Oxford Nanopore Technologies’s MinION. We picked late embryonic and adult stages because between these stages the heart undergoes structural and functional changes important for cardiac output and contractility relevant to TPM1 function.

We obtained ~310’000-740’000 reads spanning 3000-4300 unique mRNAs, 190-413 of which had a coverage of greater than 100 reads that were mapped to the rn6 genome using Minimap2 (Li 2018). We obtained average of 200,000 reads mapped to *Tpm1.* Reads that were mapped to *Tpm1* are illustrated in Figure 1A. In E20 rat hearts, we identified *Tpm1* isoforms generated via AS of exons 1a/1b, 2a/2b, 6a/6b, 9a, 9b and 9d (Figure 1A, top panel), consistent with previous findings that these mutually exclusive exons are alternatively spliced (Gunning et al. 2005b; Moraczewska et al. 1999). Strikingly, at this developmental stage *Tpm1* transcripts displayed two distinct 3’ ends defined by exon usage of either exon 9b or 9d that contain both 3’UTR and coding region (Figure 1A, top panel). There were also transcripts that ended with exon 9a (Figure 1A, 1B and 1C). The differences in the 3’end of *Tpm1* was generated via AS of terminal exons 9a-9b and 9d (Figure 1A). Inclusion of exon 9b resulted in short *Tpm1* isoforms whereas inclusion of exon 9d generated long *Tpm1* isoforms (Figure 1A, top panel). It has been previously shown that *Tpm1* isoforms that contain exon 9a-9b are primarily expressed in striated muscle (muscle-specific isoform), and that contain exon 9d are expressed in smooth muscle and other cell types (non-striated muscle isoform, in short non-muscle) (Gunning et al. 2005b; Moraczewska et al. 1999). Cardiac output increases and contractions become more coordinated at adult stages in comparison to embryonic stages. Consistent with this, muscle specific *Tpm1* isoforms that end with exon 9b were predominantly expressed at adult stages. On the other hand, non-muscle *Tpm1* isoforms that end with exon 9d (Figure 1A, bottom panel) were present in embryonic hearts but were dramatically decreased in adult hearts.

**Figure 1.**
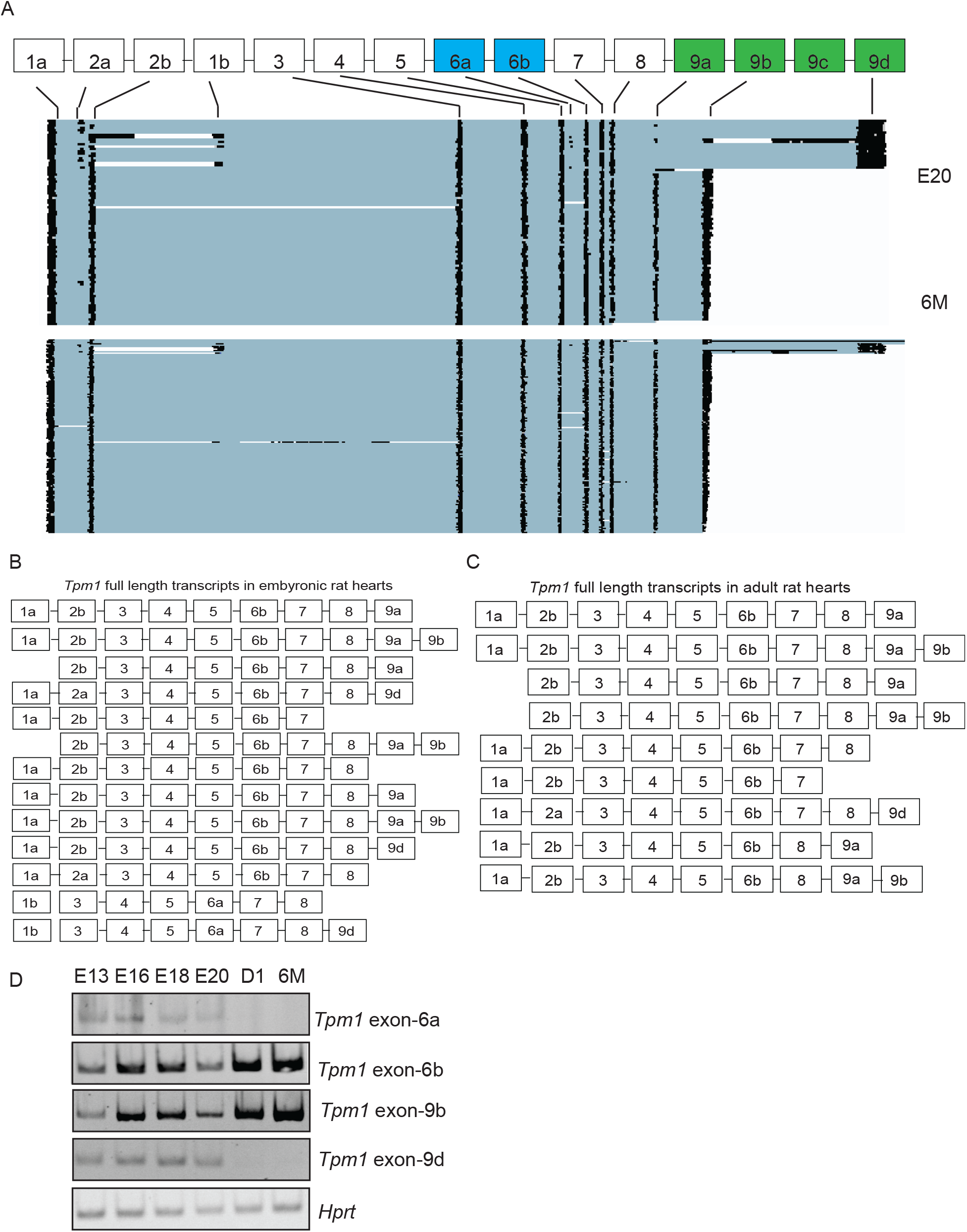
Identification of full length *Tpm1* isoforms generated via alternative splicing during rat heart development using nanopore sequencing. **(A)** Representative images of nanopore sequencing reads mapped to *Tpm1* gene in rat hearts at different developmental stages: embryonic day 20 (E20) and 6-months (6M) old (n=3). **(B)** Abundant full-length *Tpm1* isoforms in embryonic day 20 rat hearts determined by nanopore reads. **(C)** Abundant full-length *Tpm1* isoforms identified in 6 months old rat hearts by nanopore sequencing. **(D)** Relative ratio of *Tpm1* isoforms that include exon 6b vs 6a or exon 9b vs 9d in rat hearts at E13, E16, E18, E20, D1 (1-day old) and 6M.

We mapped all the exons and obtained full length *Tpm1* transcripts using the nanopore reads (Supplemental Excel File 1). In rat hearts, *Tpm1* exons 1a, 2b, 6b, 9a and 9b were more frequently used (Figure 1A, 1B and 1C). Several *Tpm1* transcripts started with exon 2b instead of exon 1a or exon 1b (Figure 1B and 1C). Interestingly, several *Tpm1* transcripts ended with exon 7 or 8 instead of exon 9a, 9d or 9d (Figure 1B and 1C). We also found unique linkages between internal and terminal exons in a given *Tpm1* transcript. For example, exon 9b was present in all muscle specific *Tpm1* transcripts that included exon 9a in rat hearts (Figure 1A, 1B and 1C). This is in agreement with predominant expression of *Tpm1* isoforms that contain both exon 9a and 9b at adult stages when the heart contracts more coordinately in comparison to the embryonic heart. Similar to exon9a-9b linkage, we found that exon 2a was almost always included in nonmuscle *Tpm1* isoforms that ended with exon 9d (Figure 1A, top panel and 1B). Exon 2a-9d containing isoforms were barely detectable in adult rat hearts (Figure 1A, bottom panel and 1C). Also, *Tpm1* isoforms that contain exon 6a were present scarcely at E20 rat hearts (Figure 1A and 1B), but they were diminished in adult rat hearts (Figure 1A and 1C). When we examined the most abundant *Tpm1* full length transcripts in E20 hearts, we noticed that alternative exons 1b, 2a, 6a and 9d were more frequently used in *Tpm1* full transcripts (Figure 1B) than that adult rat hearts (Figure 1C). These exons are associated with non-striated muscle isoforms of *Tpm1.*

To validate the nanopore sequencing data and assess expression of different *Tpm1* isoforms in embryonic rat hearts at E13, E16, E18, E20, at 1-day old (D1) vs 6-months old stages, we designed primers to determine the inclusion of exons 6a, 6b, 9b and 9d via RT-qPCR (Table S1). Consistent with the nanopore sequencing data, the expression of *Tpm1* transcripts that include muscle enriched exons 6b and 9b gradually increased dramatically after birth (Figure 1D) and expression of non-muscle exons 6a and 9d containing Tpm1 transcripts were downregulated after birth (Figure 1D). Exons 9a and 9b are important determinants for actin binding affinity of TPM1 and interactions with troponin complex in the presence or absence of Ca^2+^ (Hammell and HitchcockDeGregori 1996; Moraczewska et al. 1999). The exon 6a is required for actin binding and replacing exon 6b with exon 6a increases TPM1 actin binding affinity (Hammell and HitchcockDeGregori 1997). The differences in inclusion of internal exon 6a, exon 2a and terminal exons 9a-9b and 9d in *Tpm1* isoforms expressed in rat hearts correlates with the actin binding activity of *Tpm1* during muscle contraction at different developmental stages.

### The RNA binding protein RBFOX2 controls AS of developmentally regulated exons *of Tpm1*

RBFOX2 is an AS regulator abundantly expressed in skeletal and heart muscle (Nutter et al. 2017; Nutter et al. 2016; Singh et al. 2018b; Singh et al. 2014). RBFOX2 binding sites are enriched in introns flanking alternative exons that are regulated during postnatal mouse hearts (Kalsotra et al. 2008). To determine the role of RBFOX2 in developmentally regulated AS of *Tpm1,* we depleted RBFOX2 in embryonic rat heart derived H9c2 cells (Figure 2A). We performed RT-qPCR to validate AS of mutually exclusive exons 6b vs 6a. *Tpm1* isoforms including exon 6a were dramatically increased upon RBFOX2 depletion (Figure 2B). These results show that RBFOX2 KD altered AS and of *Tpm1* exon 6a, which is present in non-muscle *Tpm1* isoforms (Figure 1A) (Gunning et al. 2005b; Moraczewska et al. 1999).

**Figure 2.**
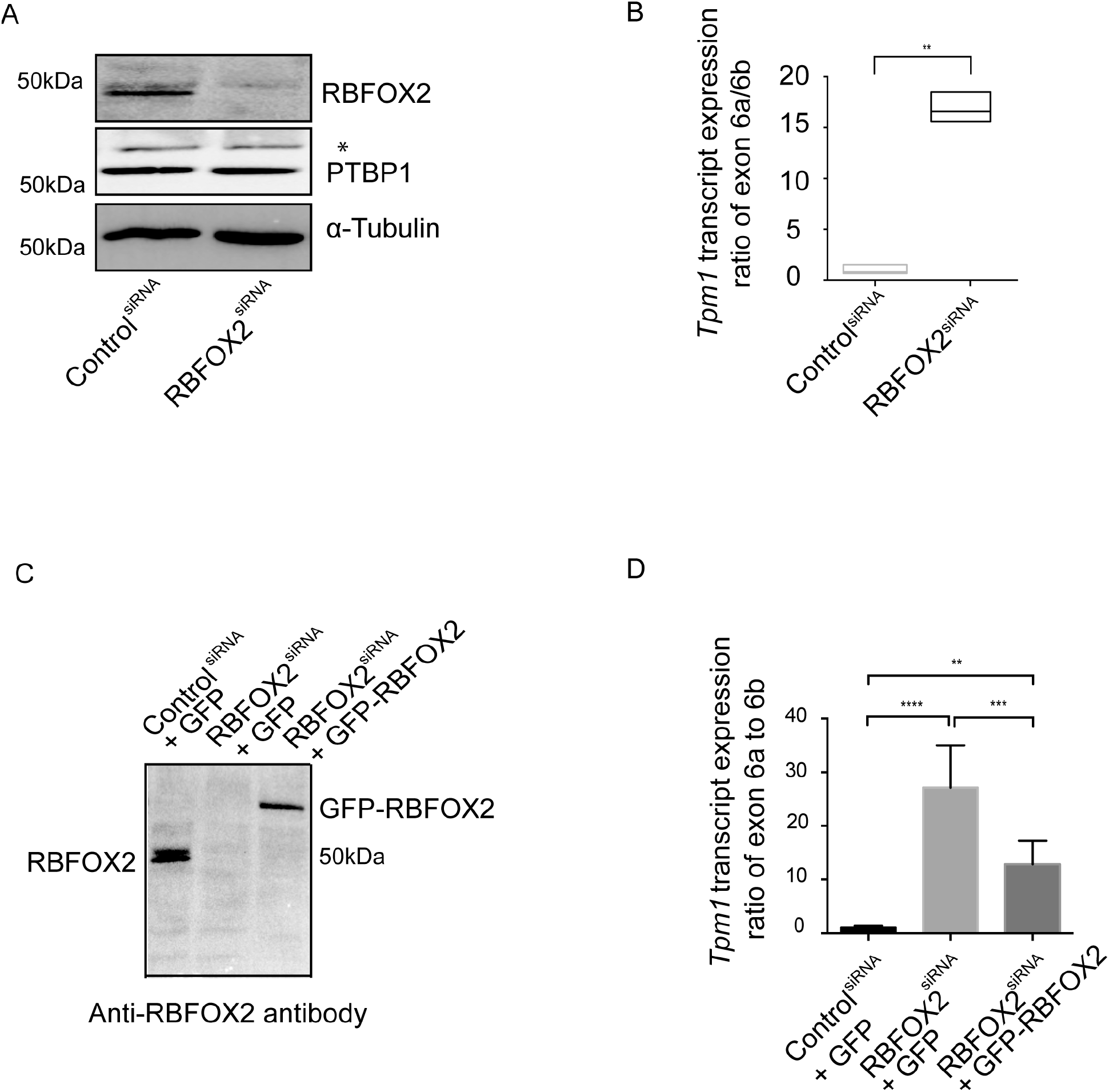
RBFOX2 regulates AS of rat Tropomyosin 1 *(Tpm1)* exon 6a. **(A)** Representative Western blot images of RBFOX2 and PTBP1 in control and RBFOX2 depleted H9c2 cells. α-tubulin was used as a loading control. **(B)** The ratio of expression levels of *Tpm1* exons 6a vs 6b in control and RBFOX2 depleted H9c2 cells determined by RT-qPCR. Expression levels of *Tpm1* exon 6a to 6b in control cells were normalized to 1. Data represent means ± SD. Statistical significance was calculated using t-test to compare two different groups in three independent experiments (n=3). ** p=0.013. GFP-REBFOX2 rescue experiments: **(C)** Western blot analysis of endogenous or GFP-tagged RBFOX2 protein in (lane1) scrambled siRNA treated, (lane 2) RBFOX2 siRNA treated, (lane 3) RBFOX2 siRNA treated H9c2 cells ectopically expressing of GFP or GFP-RBFOX2 using anti-RBFOX2 antibody. **(D)** The expression level ratios of *Tpm1* transcripts containing exons 6a vs 6b in control, RBFOX2 depleted, or RBFOX2 depleted GFP or GFP-RBFOX2 expressing cells. Expression ratios in control (1) cells were normalized to 1. Data represent means ± SD. Statistical significance was calculated using one-way ANOVA to compare three different groups in three independent experiments (n=3). P-value for Control^siRNA^+GFP vs. RBFOX2^siRNA^+GFP is p<0.000001; for RBFOX2^siRNA^+ GFP vs. RBFOX2^siRNA^+ GFP-RBFOX2 is p=0.000683; for Control^siRNA^+GFP vs. RBFOX2^siRNA^+ GFP-RBFOX2 is p=0.003409.

The RNA binding protein PTBP1 has been shown to regulate *Tpm1* AS (Gooding et al. 2013a; Lin and Tarn 2005; Llorian et al. 2010; Mullen et al. 1991; Xue et al. 2009). RBFOX2 has been shown to regulate AS and mRNA levels of PTBP family member (Jangi et al. 2014). To rule out the possibility that the effect of RBFOX2 on AS of *Tpm1* was mediated via changes in PTBP1, we examined PTBP1 protein levels in control and RBFOX2 depleted H9c2 cells. RBFOX2 KD did not affect PTBP1 protein levels (Figure 2A).

### Ectopic expression of RBFOX2 partially rescues developmentally regulated AS of *Tpm1* in RBFOX2 depleted cells

We next tested if ectopic expression of RBFOX2 can rescue *Tpm1* AS changes in RBFOX2 depleted cells. We expressed GFP tagged RBFOX2 in RBFOX2-depleted H9c2 cells and found that GFP-RBFOX2 protein was expressed at low levels similar to the endogenous RBFOX2 levels in RBFOX2 KD cells (Figure 2C, lane 1 vs 3). We tested AS of *Tpm1* exons 6a/6b in RBFOX2 KD cells expressing either GFP or GFP-RBFOX2. Expression of GFP-RBFOX2 partially rescued AS changes of *Tpm1* exon 6a/6b (Figure 2D), suggesting that RBFOX2 is a regulator of AS of developmentally regulated *Tpm1* alternative exon 6a.

### RBFOX2 and PTBP1 antagonistically control developmentally regulated AS of *Tpm1*

The RNA binding protein PTBP1 is a known regulator of *Tpm1* AS of mutually exclusive exons 2a/2b (Gooding et al. 2013a; Mullen et al. 1991) and exons 6a/6b (Xue et al. 2009) and terminal exons (Lin and Tarn 2005; Llorian et al. 2010). AS of these *Tpm1* exons are also developmentally regulated. We found that RBFOX2 also regulates AS of exons 6a/6b. To determine where these RNA binding proteins bind with respect to the developmentally regulated alternative exons of *Tpm1,* we examined the enhanced crosslinking immunoprecipitation RNA-seq (eCLIP) data for RBFOX2 and PTBP1 from ENCODE (Consortium 2012; Davis et al. 2018). We found binding clusters for both PTBP1 and RBFOX2 mapped in or near *Tpm1* exons 6a in both eCLIP experiments, suggesting that these RNA binding proteins may regulate *Tpm1* AS antagonistically (Figure 3A).

**Figure 3.**
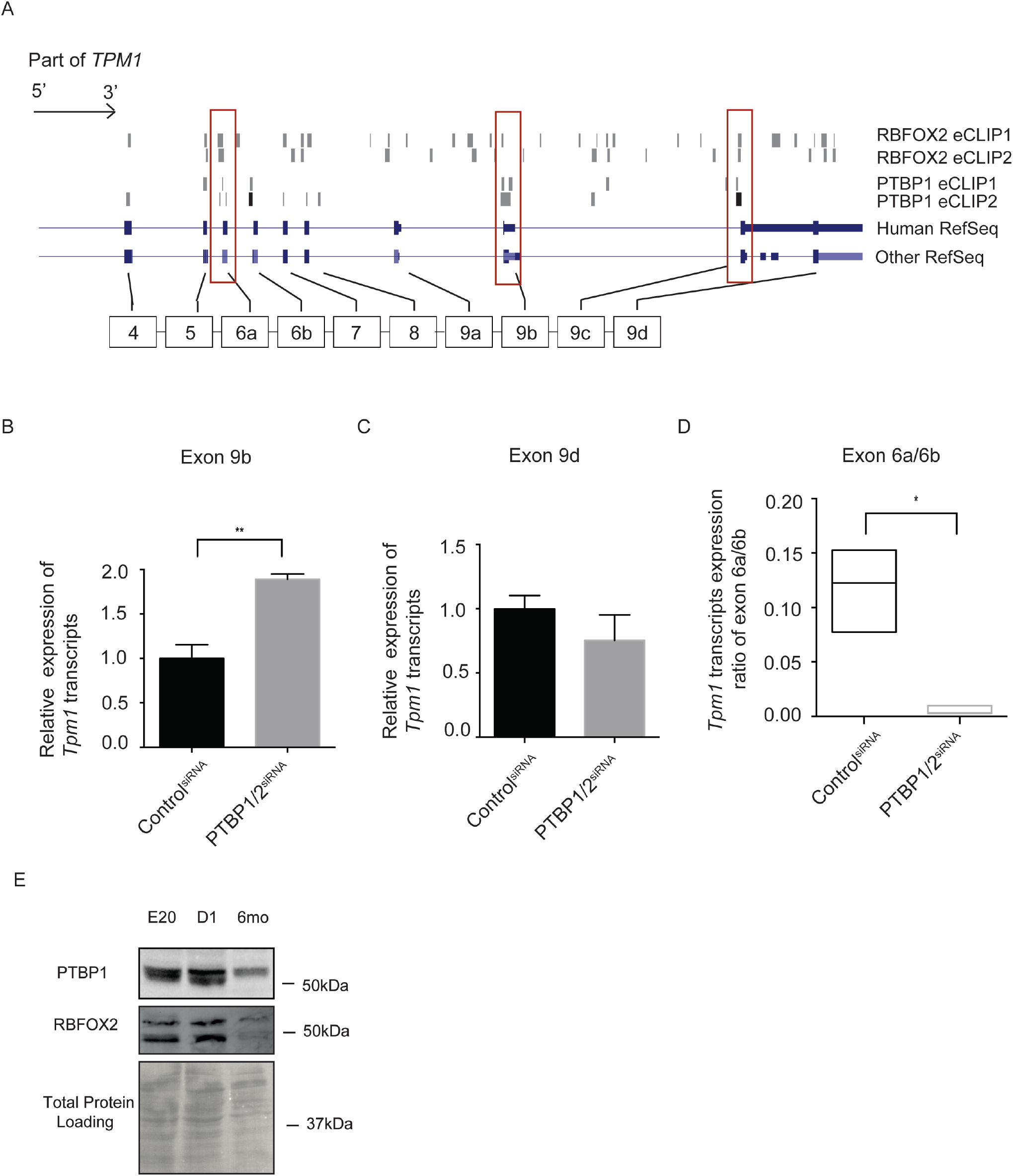
PTBP controls developmentally regulated AS of *Tpm1* antagonistically to RBFOX2. **(A)** RBFOX2 eCLIP and PTBP1 eCLIP-seq reads mapped to the human *Tpm1* gene. **(B)** Expression levels of *Tpm1* transcripts containing exon 9b (muscle) in H9c2 cells treated with control or PTBP1/2 siRNA. mRNA levels in control cells were normalized to 1. Statistical significance was calculated using t-test to compare two different groups in three independent experiments (n=3). P-value is represented as ** p= 0.0050. **(C)** Expression levels of *Tpm1* transcripts containing exon 9d (non-muscle) in H9c2 cells treated with control or PTBP1/2 siRNA. mRNA levels in control cells were normalized to 1. **(D)** Expression of *Tpm1* transcripts containing exon 6a vs 6b in H9c2 cells treated with control or PTBP1/2 siRNA. Data represent means ± SD. Expression ratio of *Tpm1* exon 6a to 6b in control cells were normalized to 1. Statistical significance was calculated using t-test to compare two different groups in three independent experiments (n=3). P-value is represented as * p= 0.0347. **(E)** Western blot analysis of PTBP1 and RBFOX2 in rat hearts at different embryonic and postnatal stages. Even protein loading was monitored by Ponceau S stain of the membrane.

PTBP1 binding sites were present in both eCLIP experiments near *Tpm1* exons 6a, 6b, and 9b (Figure 3A). To validate the regulation of these exons by PTBP, we knocked down PTBP proteins using PTBP1/2 siRNAs in embryonic rat heart derived H9c2 cells. PTBP depletion led to downregulation of muscle specific *Tpm1* transcripts that end with exon 9b (Figure 3B). There was no significant change in non-muscle *Tpm1* isoforms that end with exon 9d in PTBP1 KD cells (Figure 3C), consistent with lack of PTBP1 binding sites within or near this exon in *Tpm1* pre-mRNA (Figure 3A).

Inclusion of *Tpm1* exon 6a was inhibited in PTBP depleted H9c2 cells (Figure 3D). This was contrary to what was observed in RBFOX2 depleted H9c2 cells (Figure 2B and 2D). These findings indicate that RBFOX2 and PTBP1 antagonistically control developmentally regulated AS of *Tpm1* exon 6a, contributing to the generation of *Tpm1* isoforms with different actin binding capabilities.

To better understand how *Tpm1* muscle specific isoforms are regulated by RBFOX2 and PTBP1 during rat heart development, we checked RBFOX2 and PTBP1 protein levels during rat heart development. Both PTBP1 and RBFOX2 protein levels were abundant at embryonic stages and 1-day old pups (Figure 3E) but levels went down in adult stages. Because PTBP1 is a repressor of muscle specific *Tpm1* isoforms, low levels of PTBP1 in adult rat hearts (Figure 3E) correlated well with predominant expression of *Tpm1* muscle specific isoforms in adult hearts (Figure 1A, bottom panel and 1C).

### Developmentally regulated AS patterns of TPM2 and TPM3

There are four family members of TPM in vertebrates namely TPM1, TPM2, TPM3 and TPM4. TPM1 and TPM2 are predominantly expressed in muscle and are involved in contraction (Dube et al. 2014; Jagatheesan et al. 2010; Wieczorek et al. 2008; Yin et al. 2015), whereas TPM3 and TPM4 are enriched in non-muscle cells supporting actin cytoskeleton (Bailey 1948; Dube et al. 2014; Gunning et al. 2008; Gunning et al. 2005a; Helfman et al. 1986; Lin et al. 2008; Marston and Redwood 1993; Perry 2001; Rao et al. 2012; Yin et al. 2015). To determine whether AS of other *TPM* genes were also regulated during rat heart development, we examined our nanopore sequencing data. *Tpm2* and *Tpm3* displayed AS transitions between embryonic and adult stages in rat hearts as well as changes in its expression levels (Figure 4A and 4B top vs bottom panels). *Tpm2* isoforms that end with exon 9d were predominant in adult rat hearts (Figure 4A). Similar to *Tpm1, Tpm2* exons 6a/6b and 9b/9d differentially spliced during rat heart development (Figure 4A vs Figure 1A, 1B and 1C). *Tpm3* exons 1a/1b, exons 6a/6b and exons 9a/9b were also developmentally regulated. Interestingly, AS of *Tpm4,* which is enriched in nonmuscle cells, was not regulated via AS during rat heart development (Figure 4C).

**Figure 4.**
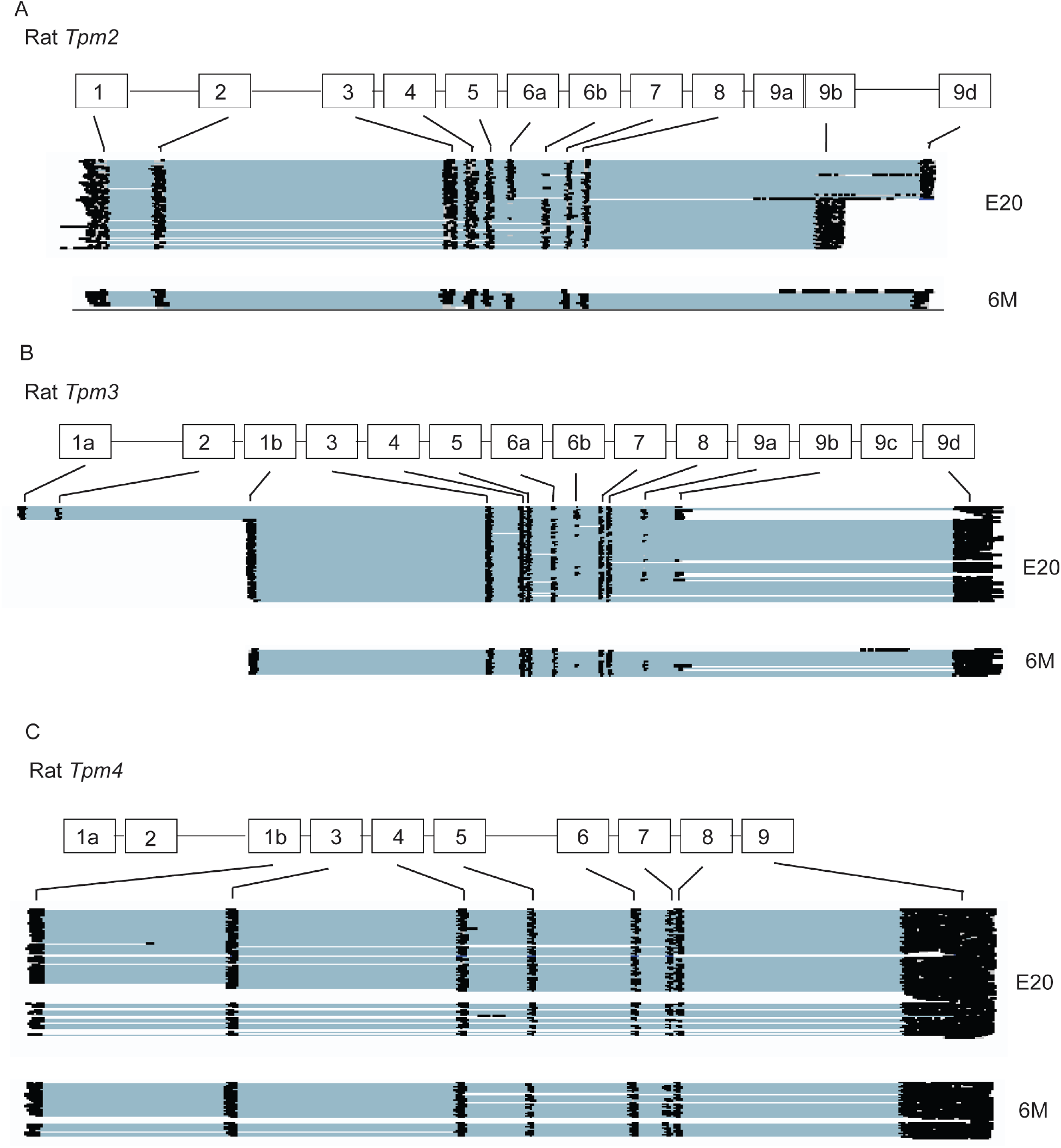
Full-length isoforms of *Tpm2* **(A)**, *Tpm3* **(B)** and *Tpm4* **(C)** identified by nanopore sequencing in embryonic day 20 (E20) and 6-months old (6M) rat hearts.

## DISCUSSION

*Tpm1* is an essential gene required for organization of the myofibril (Thomas et al. 2010), myocardial contraction (Wolska and Wieczorek 2003), and heart development (England et al. 2017). *Tpm1* has 15 exons, in which internal exons 1a/1b, 2a/2b and 6a/6b and terminal exons 9a, 9b, 9c and 9d are alternatively spliced (Crawford and Patton 2006; Dye et al. 1998; Geeves et al. 2015; Gooding et al. 2013b; Lin and Tarn 2005; Lin et al. 2008; Mullen et al. 1991). AS of *Tpm1* isoforms are tissue- and development-specific and exhibit distinct physiological functions (Gunning et al. 2015; Lin et al. 2008; Perry 2001; Schevzov et al. 2011) including cytoskeleton support for almost every eukaryotic cells and muscle contraction for striated muscle cells (Dube et al. 2014; Jagatheesan et al. 2010; Wieczorek et al. 2008; Yin et al. 2015). Full length *Tpm1* isoforms that are generated via extensive AS regulation in different cell types and at different developmental stages have not been obvious using short read RNA sequencing methods. It is because short read RNA sequencing is unable to provide direct information about how different exons are connected and incorporated into full length transcripts to generate different *Tpm1* isoforms. In addition, bias is generated from fragmentation during library preparation. Using nanopore long read sequencing by MinION, we were able to detect full length isoforms of abundant cardiac genes including tropomyosin family members without extensive RNA manipulation and gene specific PCR amplification.

Nanopore sequencing identified full length *Tpm* isoforms with different internal and terminal exon combinations in rat hearts that are regulated during development. We defined striated muscle vs non-striated muscle *Tpm1* isoforms based on the specific exon combinations and internal-terminal exon linkages in embryonic vs adult hearts. We also observed that striated muscle specific *Tpm1* isoforms were abundantly expressed at both embryonic and adult stages, but they became the predominant isoform in adult hearts. Embryonic rat hearts displayed more non-striated muscle *Tpm1* isoforms in comparison to adult rat hearts. Loss of non-muscle specific *Tpm1* isoforms in adult rats is in agreement with the increased muscle contraction capability of adult hearts in comparison to embryonic hearts.

We identified RBFOX2 as a regulator of developmentally regulated AS of *Tpm1,* consistent with a recent study has identified a splicing change in TPM1 in mouse embryos in which RBFOX2 was conditionally ablated in neural crest cells (Cibi et al. 2019). Our work provide evidence that changes in *Tpm1* AS may contribute to heart and muscle defects observed in RBFOX2 loss of function in human heart diseases and in experimental animal models (Sebastian et al. 2013),(Singh et al. 2018a),(Gallagher et al. 2011), (Homsy et al. 2015; McKean et al. 2016; Verma et al. 2016),(Wei et al. 2015),(Nutter et al. 2017; Nutter et al. 2016).

Using nanopore sequencing we validated previous findings that that AS of exons 6a/6b of *Tpm1* are developmentally regulated in rat hearts and identified new full length transcripts of *Tpm1* regulated during rat heart development. Here we showed that AS of exons 6a vs 6b were controlled antagonistically by RBFOX2 and PTBP. The exon 6a/6b is required for cooperative actin binding (Hammell and HitchcockDeGregori 1997). Therefore, the dynamic regulation of exon 6a via RBFOX2 and PTBP during cardiac development is critical to ensure selective *Tpm1* isoforms expressed in equilibrium to exert specific actin binding activity during different states of muscle contraction.

Our results support the idea that RBFOX2 is a repressor of *Tpm1* exon 6a inclusion. Conversely, PTBP1 is an activator of exon 6a, consistent with the previous reports (Lin and Tarn 2005; Llorian et al. 2010; Xue et al. 2009). The expression levels of both PTBP1 and RBFOX2 were high at embryonic stages but both were decreased at adult stages in rat hearts. While the predominant expression of muscle specific *Tpm1* isoforms at adult stages was consistent with downregulation of PTBP1, the abundance of muscle specific *Tpm1* isoforms at embryonic stages was in agreement with high levels of RBFOX2. The interplay between these RNA binding proteins in regulating *Tpm1* AS correlates well with their different roles as repressors or activators of exon inclusion.

It is quite common that cardiac structural genes undergo cooperative AS regulation that determine their specific functions and expression profiles during cardiac development. Mutations in these structural genes including *TPM1* are linked to human heart diseases. Our results using nanopore sequencing provide as an efficient way to reveal full length isoforms of cardiac structural genes and their regulation during heart development. In addition, our work may pave the way for future studies to determine the functional consequences of non-coding mutations on post-transcriptional regulation of cardiac structural genes using nanopore sequencing.

## MATERIALS AND METHODS

### Cell culture

H9c2 cells (ATCC CRL-1446) were cultured and maintained in Dulbecco’s modified Eagle’s medium (DMEM) (ATCC 30-2002), supplemented with 10% fetal bovine serum (FBS, ATCC 30-2020) and 100 units/ml penicillin and streptomycin (Thermofisher Scientific 15140122).

### Transfections

For siRNA-KD experiments, H9c2 cells were seeded at 10^6^ cells per 100mm dish and transfected with 20nM scrambled siRNA (Invitrogen AM4611), *Rbfox2* siRNA (Invitrogen siRNA ID# s96620) or PTBP siRNAs (Qiagen cat# SI02649206 and SI04255146) using Lipofectamine RNAiMAX (Thermofisher Scientific). Cells were harvested 72 hours posttransfection for RNA or protein extraction. For rescue experiments, 3X10^6^ H9c2 cells were transfected with eGFP (Sigma-Aldrich), human GFP-RBFOX2 (transcript variant 3) (Addgene, plasmid #63086) or empty vector (pcDNA 5) together with scrambled or *Rbfox2* specific siRNAs using Neon Nucleofection System (Thermofisher Scientific) as described previously (Verma et al. 2013). RNA was harvested 48 hours post-transfection.

### Nanopore sequencing with MinION

RNA was extracted from cells using TRIzol (Invitrogen 15596-018) by following the manufacturer’s protocol. For nanopore sequencing, three sets of E20 and three sets of 6M rat heart RNA (Zyagen) were used. Total cellular RNA was first poly(A) enriched (New England Biolabs S1550S) and then amplified using oligo-dT primers and template switching oligos using Oxford Nanopore Technologies (ONT) cDNA-PCR sequencing kit (PCS108) as described by the manufacturer. Samples were multiplexed using ONT barcodes. Pooled samples were sequencing on R9.4 flowcells for 36. Reads were demultiplexed and base-called using Albacore and mapped to the rat genome (rn6) using the −splice function of minimap2 as described previously (Li 2016). Analysis pipeline is shown in Supplemental Figure 1.

### RT-qPCR

Total RNA from cells and rat hearts (purchased from Zyagen) at E13 (pooled), E16 (pooled), E18 (pooled), E20 (pooled), post-natal day1 (D1) (pooled), and 6M (pooled) stages were extracted using Trizol. 2μg of total RNA was used for cDNA synthesis using AMV reverse transcriptase (15 units/μg, Life Biosciences). For RT-qPCR, master mix was set up by mixing 5 μl of cDNA, 3 μl of H_2_O, 2 μl of PCR gene specific primer (10X conc) (Table S1) and 10 μl of master mix (Roche 04707516001) in 20 μl reaction. The RT-qPCR was conducted using LightCycler 480 Instrument (Roche) using the following conditions: 95 °C 10 s; 62 °C 15 s; 72 °C 10s for 40 cycles. Melting curve was obtained to ensure single product. ΔCt method was adopted for quantification. Semi-quantitative RT-PCR instead of RT-qPCR was used for determining *Tpm1* short and *Tpm1* long transcript levels due to the alternative splicing of exon 9a that generates two different sized DNA bands after amplification. 2μg of total RNA was used for cDNA synthesis using AMV reverse transcriptase (15 units/μg, Life Biosciences). PCR was performed using 5μl of cDNA, 25μM dNTPs, 100ng of each gene specific forward and reverse primer and 0.2μl of Biolase Taq polymerase (Bioline) in a 20μL reaction. The amplified products were analyzed on 5% acrylamide gel. *Hprt* was used as an internal control for RT-PCR quantification.

### Western Blot

The membrane was blocked with 5% dry fat-free milk solution in PBS containing 0.1% Tween (PBST) at RT for 1 hour and then incubated with indicated primary antibodies overnight at 4°C. The membrane was washed with PBST for 15 minutes three times and incubated with HRP-conjugated secondary antibody for 1 hour at RT followed by three washes using PBST. Immobilon Western chemiluminescent (Millipore WBKLS0500) kit was used to detect HRP activity of the secondary antibody. The membrane was then imaged using ChemiDoc Touch imaging system (Bio-rad). Image J software was used for band intensity quantification. Primary antibodies used for this study are as follows: TPM1 (1:1000, Cell Signaling, D12H4), RBFOX2 (1:1000, Abcam, ab57154), PTBP1 (1:5000, a gift from Dr. Mariano Garcia-Blanco) and α-tubulin (1:20000, Sigma-Aldrich, T6074).

## DATA AVAILABILITY

Nanopore sequencing data was deposited to NCBI SRA database with project number PRJNA517125.

## ACKNOWLEDGEMENTS, FUNDING AND CONFLICT OF INTEREST

This work was supported, in part, UTMB Department of Biochemistry and Molecular Biology Bridging funds; and a grant from the National Institutes of Health/ National Heart Lung Blood Institute [1R01HL135031], a grant from CPRIT [RP190556] and a grant from American Heart Association [20TPA35490206] to M.N.K-M. The contents of the manuscript are solely the responsibility of the authors and do not necessarily represent the official views of NHLBI of NIH. J.C. is a funded by a post-doctoral fellowship from American Heart Association [18POST3399018]. A.R. is supported by start-up funds from UTMB. We thank Dr. Garcia-Blanco for providing us PTBP1 antibody. Authors declare no conflict of interest.

## ABBREVIATIONS

AS: alternative splicing
E20: embryonic day 20
6mo: 6-months old
KD: knock down
TPM1: Tropomyosin 1

## SUPPORTING INFORMATION (FIGURES AND TABLES)

### Supplemental Figure 1.

**Figure S1.**
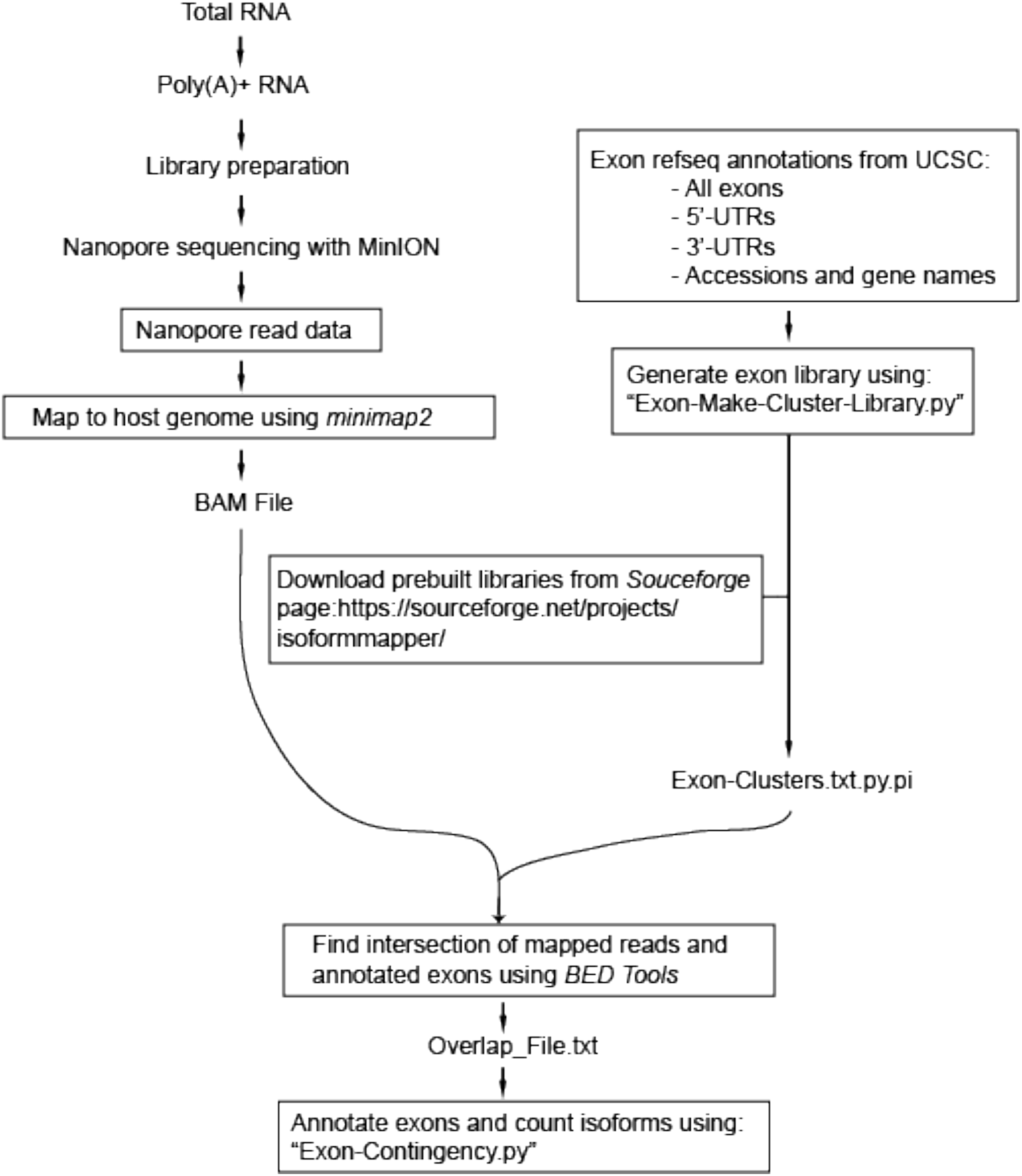
Workflow for nanopore cDNA sequencing.

### Supplemental Table 1

**Table S1.**
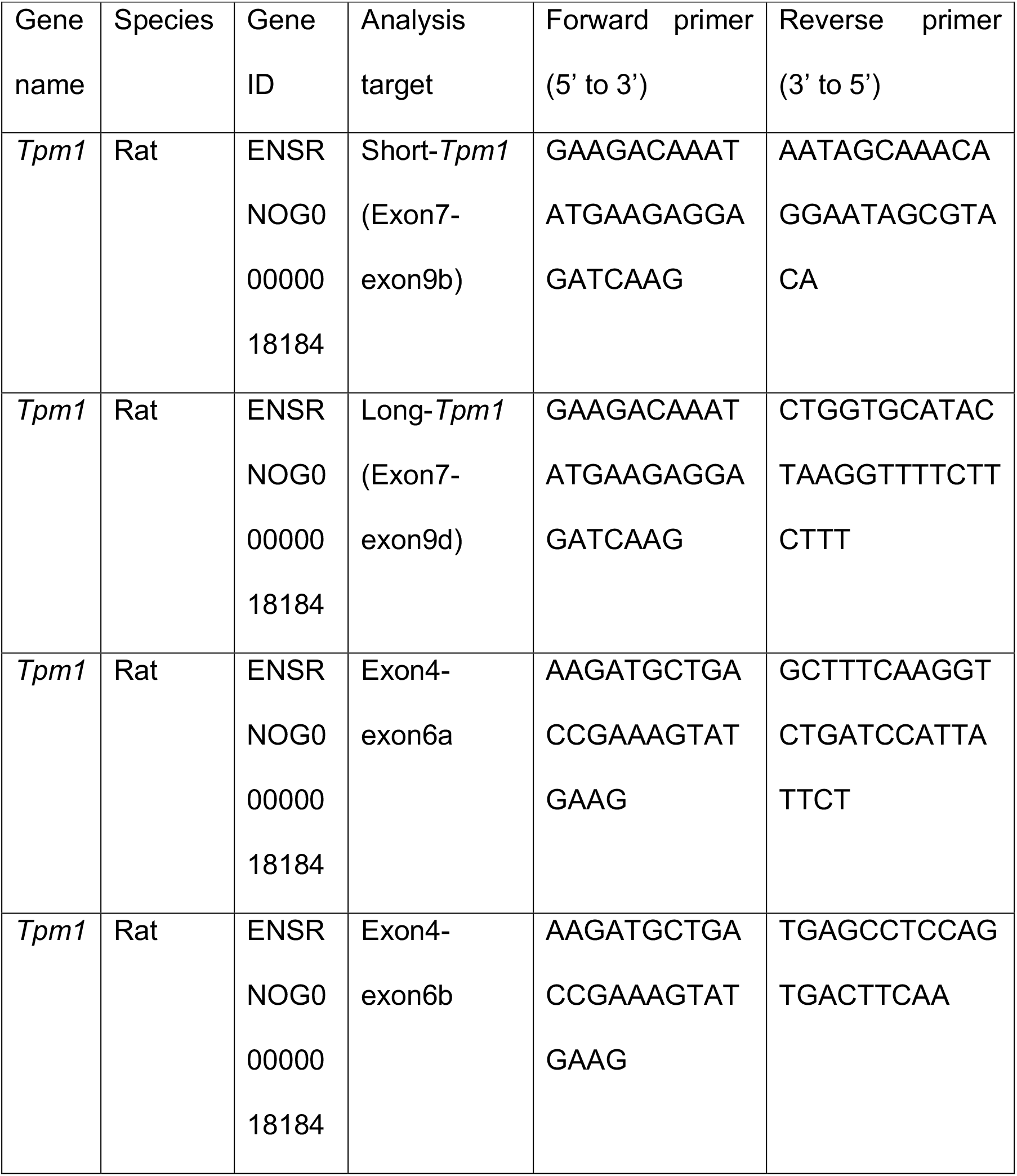
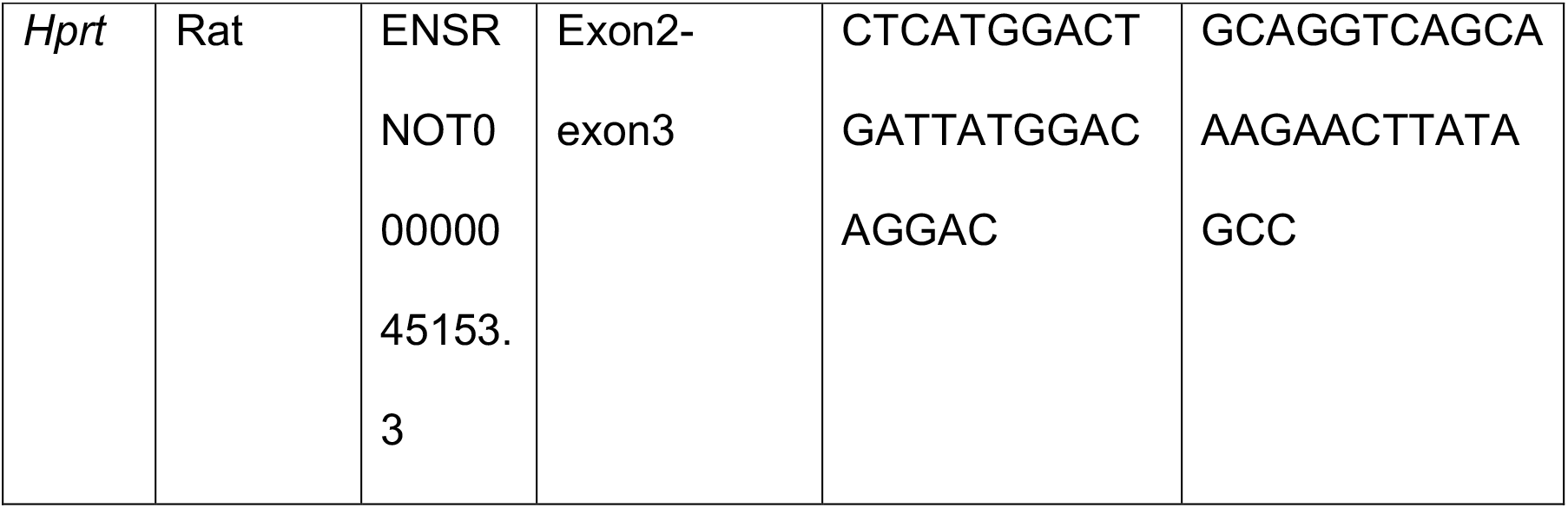
Primer list for PCR reactions.

